# Targeting Ras signaling excitability in cancer cells through combined inhibition of FAK and PI3K

**DOI:** 10.1101/2023.06.12.544386

**Authors:** Chao-Cheng Chen, Suyang Wang, Jr-Ming Yang, Chuan-Hsiang Huang

**Author notes:** Correspondence (C.-H.H.), (J.-M.Y.).

## Abstract

The Ras/PI3K/ERK signaling network is frequently mutated in various human cancers including cervical cancer and pancreatic cancer. Previous studies showed that the Ras/PI3K/ERK signaling network displays features of excitable systems including propagation of activity waves, all-or-none responses, and refractoriness. Oncogenic mutations lead to enhanced excitability of the network. A positive feedback loop between Ras, PI3K, the cytoskeleton, and FAK was identified as a driver of excitability. In this study, we investigated the effectiveness of targeting signaling excitability by inhibiting both FAK and PI3K in cervical and pancreatic cancer cells. We found that the combination of FAK and PI3K inhibitors synergistically suppressed the growth of select cervical and pancreatic cancer cell lines through increased apoptosis and decreased mitosis. In particular, FAK inhibition caused downregulation of PI3K and ERK signaling in cervical cancer but not pancreatic cancer cells. Interestingly, PI3K inhibitors activated multiple receptor tyrosine kinases (RTKs), including insulin receptor and IGF-1R in cervical cancer cells, as well as EGFR, Her2, Her3, Axl, and EphA2 in pancreatic cancer cells. Our results highlight the potential of combining FAK and PI3K inhibition for treating cervical and pancreatic cancer, although appropriate biomarkers for drug sensitivity are needed, and concurrent targeting of RTKs may be required for resistant cells.

## Introduction

The signaling network involving Ras GTPases and their downstream effectors including PI3K and MAPK/ERK pathways regulates diverse cellular processes such as cell proliferation, differentiation, apoptosis, metabolism, protein synthesis, and cell motility [1]. Activation of the network through mutations or overexpression is frequently found in human cancer [2]. For example, 10-36% of cervical cancers harbor activating mutations in PIK3CA, which codes for the catalytic subunit of PI3Kα [3–6], and KRAS mutations are found in ∼8% of non-SCC of the cervix [7](ref). The MAPK/ERK signaling pathway has also been implicated in cervical cancer pathogenesis [8,9].Clinical trials of PI3K pathway inhibitors showed promise in stabilizing the progression of metastatic cervical cancer, but the response was variable [10,11]. In pancreatic ductal adenocarcinoma (PDAC), KRAS mutations are found in more than 90% of tumors [12,13], and some hereditary pancreatic cancers have been linked to mutations that change posttranslational modification of Ras GTPases [14]. In KRAS wild-type PDAC, activation of the network often results from various mechanisms including BRAF mutation and RTK fusions [15].

Recent studies on the spatiotemporal dynamics Ras/PI3K/ERK signaling at the cellular level revealed features of excitable systems. For example, Ras and PI3K activities propagate as self-organized waves in various cell types such as Dictyostelium, neutrophils, fibroblasts, and epithelial cells [16–24], whereas ERK displays pulsatile activation in individual cells and can propagate as waves across the cell population [25–30]. Furthermore, when stimulated by growth factors or chemoattractants, activation of the signaling network displays all-or-none characteristics and refractoriness [21–23,31–33]. Our previous study demonstrated that pulsatile ERK activity is driven by localized Ras activation on protrusions, which are stochastically generated by an positive feedback loop involving Ras, PI3K, actin, and focal adhesion kinase (FAK) [23]. Chemical and mechanical stimuli may enter the feedback loop at different points to modulate the frequency of protrusions and ERK pulses. Thus, the excitability property allows the Ras signaling network to integrate different types of stimuli in the regulation of cell proliferation. We also found that oncogenic mutations induce enhanced excitability of the Ras/PI3K/ERK signaling network [22,23].

In this study, we test the hypothesis that inhibition of the molecular mechanism of Ras signaling excitability can effectively block cancer cell growth, focusing on the positive feedback loop between Ras, PI3K, actin, and FAK [23]. Due to possible redundant pathways, we reason that effective inhibition of excitability requires simultaneous blocking of multiple proteins. In support of this idea, multipoint inhibition of the same signaling pathway has been shown to be more effective than inhibiting a single node [34]. Therefore, we tested whether combined inhibition of PI3K and FAK synergistically reduces cancer cell viability since: 1) small molecule inhibitors are readily available for these kinases; 2) FAK and PI3K are hyperactivated or overexpressed in cancer such as cervical cancer and PDAC [35,36]; and 3) inhibitors of FAK or PI3K have been tested in clinical and preclinical settings with variable outcomes [37,38], but to our knowledge their combinations have not been critically evaluated. Our results showed that as a single agent, FAK inhibition caused distinct effects on PI3K and ERK signaling in different cancer cell lines, while PI3K inhibitors activated multiple receptor tyrosine kinases (RTKs), including insulin receptor and IGF-1R in cervical cancer cells, as well as EGFR, Her2, Her3, Axl, and EphA2 in pancreatic cancer cells. When combined, FAK and PI3K inhibitors synergistically suppressed the growth of select cervical and pancreatic cancer cell lines through increased apoptosis and decreased mitosis. These findings support the potential usefulness of combining FAK and PI3K inhibition for treating cervical and pancreatic cancer. Additionally, they highlight the necessity for improved biomarkers to assess drug sensitivity and the development of strategies to overcome drug resistance.

## Materials and Methods

### Cell lines

HeLa and SiHa cell lines were purchased from ATCC. Panc10.05, A6L and Panc215 cells were gifts from the lab of Laura Wood (JHU). Cells were grown at 37°C and 5% CO_2_ in DMEM high glucose medium (Gibco, #11965092) supplemented with 10% FBS (Corning Cellgro, 35-010-CV), 1 mM sodium pyruvate (Gibco, #11360070), and 1X nonessential amino acids (Gibco, #11140050). CaSki cells were grown at 37°C and 5% CO_2_ in RPMI-1640 high glucose medium (Gibco, #11965092) supplemented with 10% FBS, 1 mM sodium pyruvate, and 1X nonessential amino acids.

### Chemical reagents

Stocks of 10 mM PF-04691502 (Sigma-Aldrich, PZ0235), 5 mM ZSTK474 (Selleckchem, S1072) and 50 mM VS-6063 (defactinib; Selleckchem, S7654) were prepared by dissolving the chemicals in DMSO. All drug stocks were stored at -20°C.

### Cell viability assay

The viability of cells after drug treatment was performed by Cell Counting Kit-8 (CCK-8, Dojindo, CK04-20) according to the manufacturer’s instructions. Briefly, 3,000 cells were seeded in a 96-well plate and incubated at 37°C and 5% CO_2_ overnight before being exposed to drugs. After a designated period of incubation, the CCK-8 reagent was directly added into each well and incubated with cells for another 3h. The absorbance of each well was analyzed by an ELISA reader (800 TS absorbance reader, BioTek) equipped with a 450 nm filter.

### Analysis of synergy

Bliss synergy scores were calculated using SynergyFinder [39]. Briefly, the cell viabilities results were entered into an example table format from SynergyFinder website (https://synergyfinder.fimm.fi). The data was then uploaded to the SynergyFinder website. The four-parameter logistic regression (LL4) was chosen for the curve-fitting algorithm. The outlier detection and Bliss model were selected for calculating drug combination effect.

### RealTime-Glo MT Cell Viability Assay

Monitoring cell viability in real time was performed by the RealTime-Glo MT Cell viability assay kit (Promega, G9713) according to the manufacturer’s instructions. Briefly, after three thousand cells were plated and incubated in a 96-well white opaque plate overnight, the medium was replaced with a phenol red-free medium containing vehicle (DMSO), PF-04691502, VS-6063 or both drugs. 1000X MT Cell Viability Substrate and 1,000X NanoLuc^®^ Enzyme were diluted to 2X with medium, and then added to each well in equal volumes. The luminescence intensity was measured by a BMG LABTECH FLUOstar Omega Microplate Reader (Cary, NC, USA).

### Cell cycle, cell growth and apoptosis analysis by flow cytometry

Flow cytometry was used to examine DAPI for cell cycle analysis, Ki-67 for cell proliferation, and cleaved caspase 3 expression for apoptosis following drug treatment. Cells were fixed with 4% formaldehyde and permeabilized using ice-cold methanol. The cells were then incubated with DAPI (10 mg/mL, 1:1000), mouse anti-human Ki-67 (Cell Signaling #9449, 1:400), and rabbit anti-human cleaved caspase 3 antibodies (Cell Signaling #9661, 1:800) at room temperature for 1 hour. Anti-mouse IgG DyLight 594 (Invitrogen #35511, 1:100) and anti-rabbit IgG DyLight 488 (Invitrogen #35553, 1:100) were used to detect anti-Ki-67 and anti-cleaved caspase 3 antibodies, respectively. The experiments were carried out on a CytoFLEX Flow Cytometry (Beckman Coulter, CA, USA), and the data was analyzed with FlowJo v10 from BD Biosciences (Franklin Lakes, NJ, USA).

### Immunoblotting

The following antibodies were purchased from Cell Signaling Technology: anti-FAK (#13009, 1:1000), anti-phospho-FAK (#8556, 1:500), anti-AKT (#2920, 1:1000), anti-phospho-AKT (#4060, 1:1000), anti-ERK (#9107, 1:1000), anti-phospho-ERK (#9101, 1:1000), GAPDH (#2118, 1:2000), Donkey anti-Rabbit Alexa Fluor™ 647 (ThermoFisher, A-31573, 1:5000), and Donkey anti-Mouse Alexa FluorTM 647 (ThermoFisher, A-31571, 1:5000).

The procedure of immunoblotting has been described in our previous publication [23]. Briefly, cells were harvested and lysed in 1X RIPA buffer (Cell Signaling, #9806) containing 1X protease inhibitor cocktail (Roche, #11873580001) and 1X phosphatase inhibitor (Sigma, #P5726). Cell lysates were collected after centrifugation and then stored at -80°C if not used immediately. After mixing with sample buffer and boiling at 95°C for 5 min, proteins were separated by SDS-PAGE using 4–20% Criterion™ TGX™ Precast Midi Protein Gels (Bio-Rad, #5671094), and then transferred to the low fluorescence background PVDF membrane (Millipore, #IPFL00005) in an ice bath at 80V for 1h. Membranes were incubated with primary antibodies overnight followed by incubation with secondary antibodies in 5% BSA in TBST at room temperature for 1h. Images were taken by the Pharos Molecular Imager (BioRad) and analyzed using ImageJ. Background was subtracted from intensities of individual protein bands. Phosphoprotein levels were normalized to either GAPDH or total level of the corresponding protein.

### Phospho-RTK array

The receptor tyrosine kinase (RTK) activities in cells after drug treatment were analyzed by a Proteome Profiler Human Phospho-RTK Array Kit (R&D Systems, ARY001B) according to the manufacturer’s instructions. After exposure to drugs for the designated periods, cells were harvested and lysed by lysis buffer provided in the kit (R&D Systems, #895943). The supernatant was collected after centrifugation at 14,000xg for 5 min at 4°C. The array membrane was soaked in the blocking buffer at room temperature for 1 h and incubated with lysed samples at 4°C overnight. After washing with the wash buffer, the membrane was immersed in secondary antibodies at room temperature for 2 h. The images were acquired using the ChemiDoc™ Touch Imaging System (Bio-Rad).

## Results

### Synergy between FAK and PI3K inhibition on the growth of cervical and pancreatic cancer cells

We investigated the impact of combined PI3K and FAK inhibition on cancer cell growth. First, we assessed the IC50 values of PF-04691502 (a PI3K/mTOR dual inhibitor) and VS-6063 (defactinib, a FAK inhibitor) in suppressing the growth of three cervical cancer lines: HeLa, SiHa, and CaSki. In these cell lines, the IC50 was approximately 0.1 μM for PF-04691502 and 5 μM for VS-6063 (**Fig. S1A**). Subsequently, we treated the cells with combinations of varying concentrations of PF-04691502 and VS-6063. Evaluation of the Bliss score revealed that PF-04691502 and VS-6063 synergistically inhibited the growth of HeLa cells, but not SiHa or CaSki cells (**Fig. 1A, B**). To broaden our analysis, we performed similar experiments in two pancreatic cancer cell lines, A6L and Panc10.05. Encouragingly, we observed synergistic inhibition of growth in both cell lines when treated with PF-04691502 and VS-6063 (**Fig. 1C, D**).

**Figure 1.**
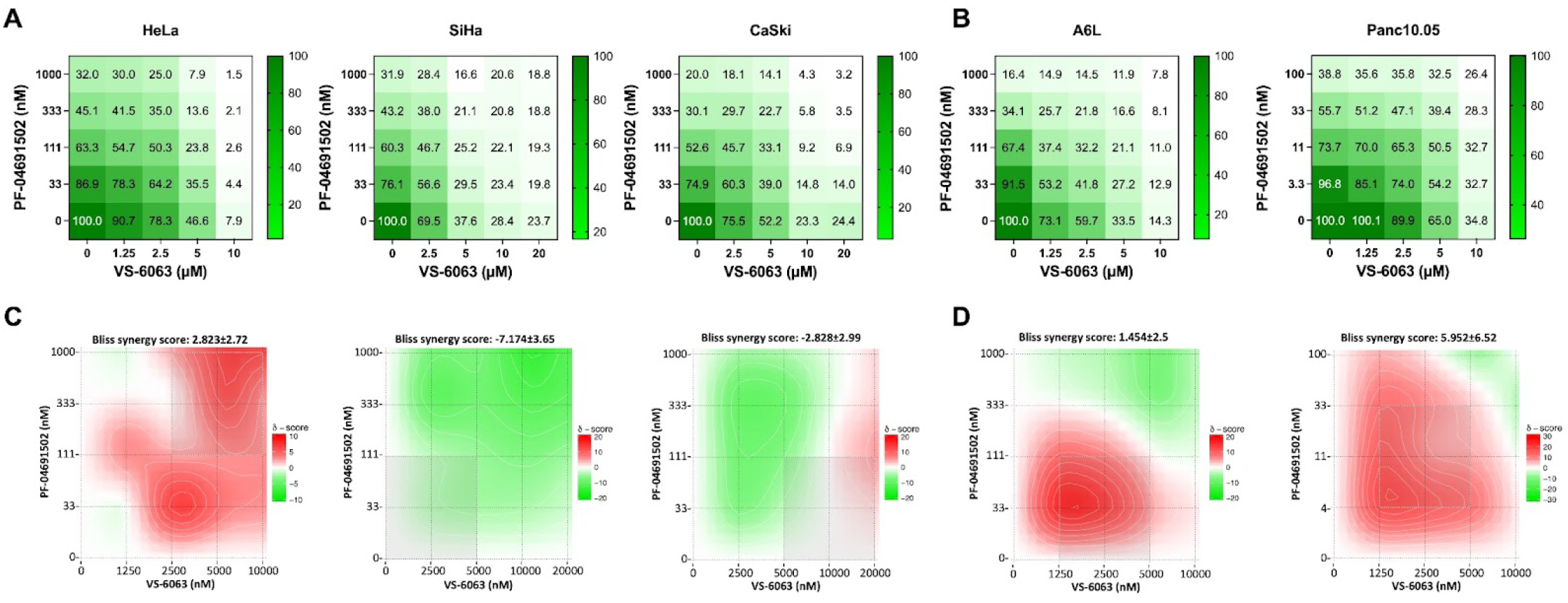
Synergy between PI3K and FAK inhibitors in cervical and pancreatic cancer cell lines. (A-B) Viability of cervical cancer cell lines (A) and pancreatic cancer cell lines (B) measured by CCK-8 assay after treatment with indicated combinations of PF-04691502 and VS-6063 for 48 hours. The numbers represent the percentage of viability compared to that of DMSO control. (C-D) Bliss synergy score for PF-04691502 and VS-6063 calculated from (A-B) using SynergyFinder [39].

To gain insight into the dynamics of cell growth in response to PI3K and FAK inhibitor combinations, we utilized the RealTime-Glo MT Cell Viability Assay to track cell behavior over a 72-hour period. For HeLa cells, both

PF-04691502 and VS-6063 exhibited cytostatic effects as individual agents but demonstrated cytocidal effects when combined (**Fig. 2A**). In contrast, the effects of inhibitors on SiHa, A6L, and Panc10.05 were mainly cytostatic. To further investigate the impact of the inhibitors on cell growth and death, we treated HeLa cells with inhibitors followed by flow cytometry analysis of Ki-67, caspase-3, and DAPI staining at 8, 12, and 24 hours after treatment (**Fig. 2B, Fig. S2**). Mitotic (M phase) cells can be identified by a high Ki-67 level and DNA content determined by DAPI staining (**Fig. 2B,C**, red boxes). VS-6063 treatment led to a significant decrease in M phase cells (**Fig. 2D**) and increase in caspase-3 (+) cells (**Fig. 2E**). Interestingly, PF-04691502 by itself had limited effects on cell cycle progression and cell death, but synergized with VS-6063 to reduce M phase cells and increase caspase-3 (+) cells (**Fig. 2D, E**). Together, these findings suggest that PF-04691502 and VS-6063 work together to increase cell death and inhibit cell cycle progression.

**Figure 2.**
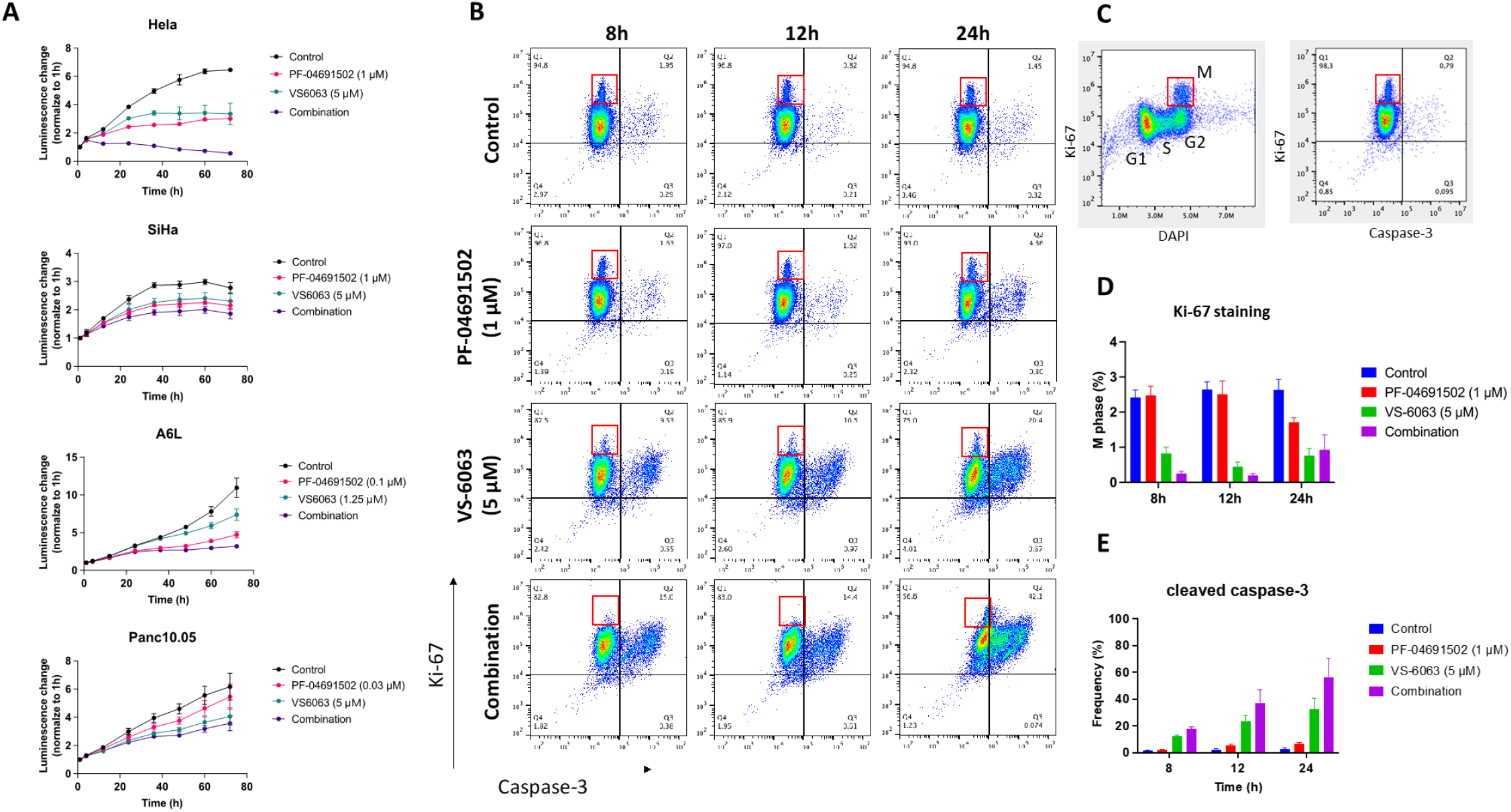
Effects of combined PI3K and FAK inhibition on cell proliferation, apoptosis, and cell cycle. (A) Proliferation of cells treated with indicated concentrations of PF-04691502 and VS-6063 tracked using the RealTime-Glo MT Cell Viability Assay. (B) Flow cytometry analysis of caspase-3 and Ki-67 staining of HeLa cells treated with indicated concentrations of PF-04691502 and VS-6063 for 8h, 12h, and 24h. (C) Example of cell cycle phase determination by Ki-67 and DAPI staining. In (B) and (C), red boxes indicate M phase cells. (D-E) Quantification of M phase (red boxes) and caspase-3 (+) cells in (B). The numbers represent mean ± SEM of n=3 experiments.

### Signaling effects of PI3K and FAK inhibitors in cervical and pancreatic cancer cells

To assess the effects of inhibitors on signaling activities, we conducted immunoblotting experiments, focusing on PI3K, ERK, and FAK pathways. Treatment with PF-04691502 led to sustained suppression of phospho-AKT in cervical cancer cell lines SiHa and CaSki (the level was undetectable in HeLa) but only transient suppression of phospho-AKT in pancreatic cancer cell lines Panc10.05, A6L, and Panc215 (**Fig. 3**). To rule out that the pAKT rebound was due to inhibitor degradation, we collected conditioned media from cells treated with the inhibitor for 0∼72 hours. When the conditioned media were added to fresh cells, phospho-AKT was inhibited, suggesting that the PI3K inhibitor retained its activity (**Fig. S3**). Thus, reactivation of AKT in the presence of PI3K inhibition was not due to loss of inhibitor potency. PF-04691502 also caused an increase in phospho-FAK over 72 hours, but the kinetics differed between cell lines (**Fig. 3**).

**Figure 3.**
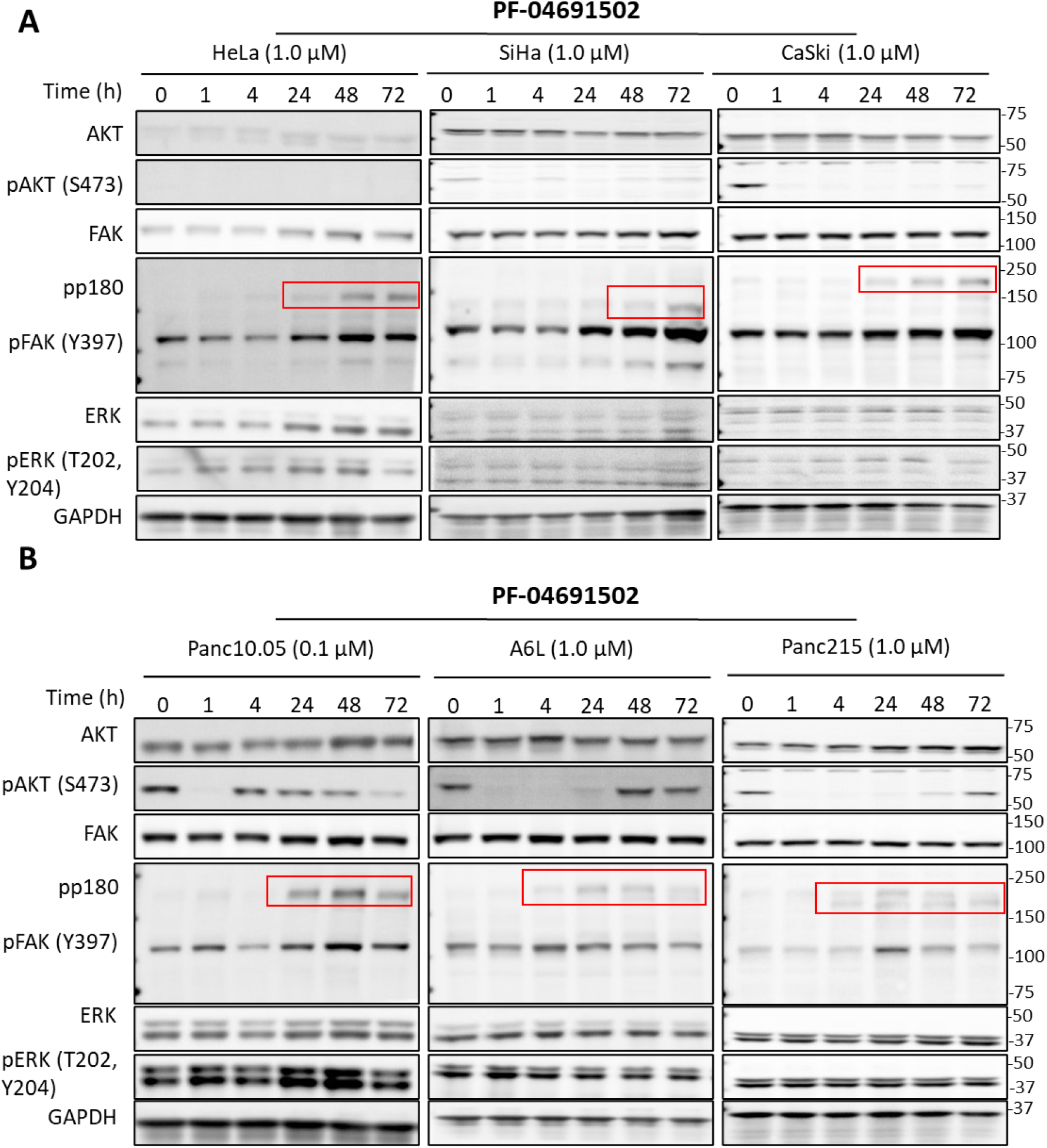
Immunoblotting of cervical (A) and pancreatic (B) cancer cells treated with PF-04691502. Cells were treated with PF-04691502 for different incubation periods immunoblotted for total and phosphorylated forms of AKT, FAK, and ERK. GAPDH was used as a loading control.

In contrast, VS-6063 caused sustained reduction in phospho-FAK in both cervical and pancreatic cell lines (**Fig. S4**). VS-6063 also caused reduced phospho-ERK in cervical cancer cell lines (**Fig. S4A**). Interestingly, VS-6063 caused increased phospho-ERK levels in pancreatic cell lines (**Fig. S4B**).

When HeLa cells were exposed to a combination of PF-04691502 and VS-6063, we observed a significant decrease in both phospho-ERK and phospho-FAK due to cell death within 24 hours (**Fig. 4A**). In contrast, the combination of PF-04691502 and VS-6063 caused a significant increase in phospho-ERK in both Panc10.05 (**Fig. 4B**) and A6L (**Fig. S5**). Taken together, these results suggest that PI3K and FAK inhibitors have distinct effects on cervical and pancreatic cancer cell lines, notably the opposite effect on phospho-ERK. However, it is crucial to note that the magnitude and kinetics of these changes exhibited variations across different experiments and cancer cell lines. These findings signify the existence of complex and non-linear interactions within the RTK signaling network, which are further influenced by the specific characteristics of each cell line.

**Figure 4.**
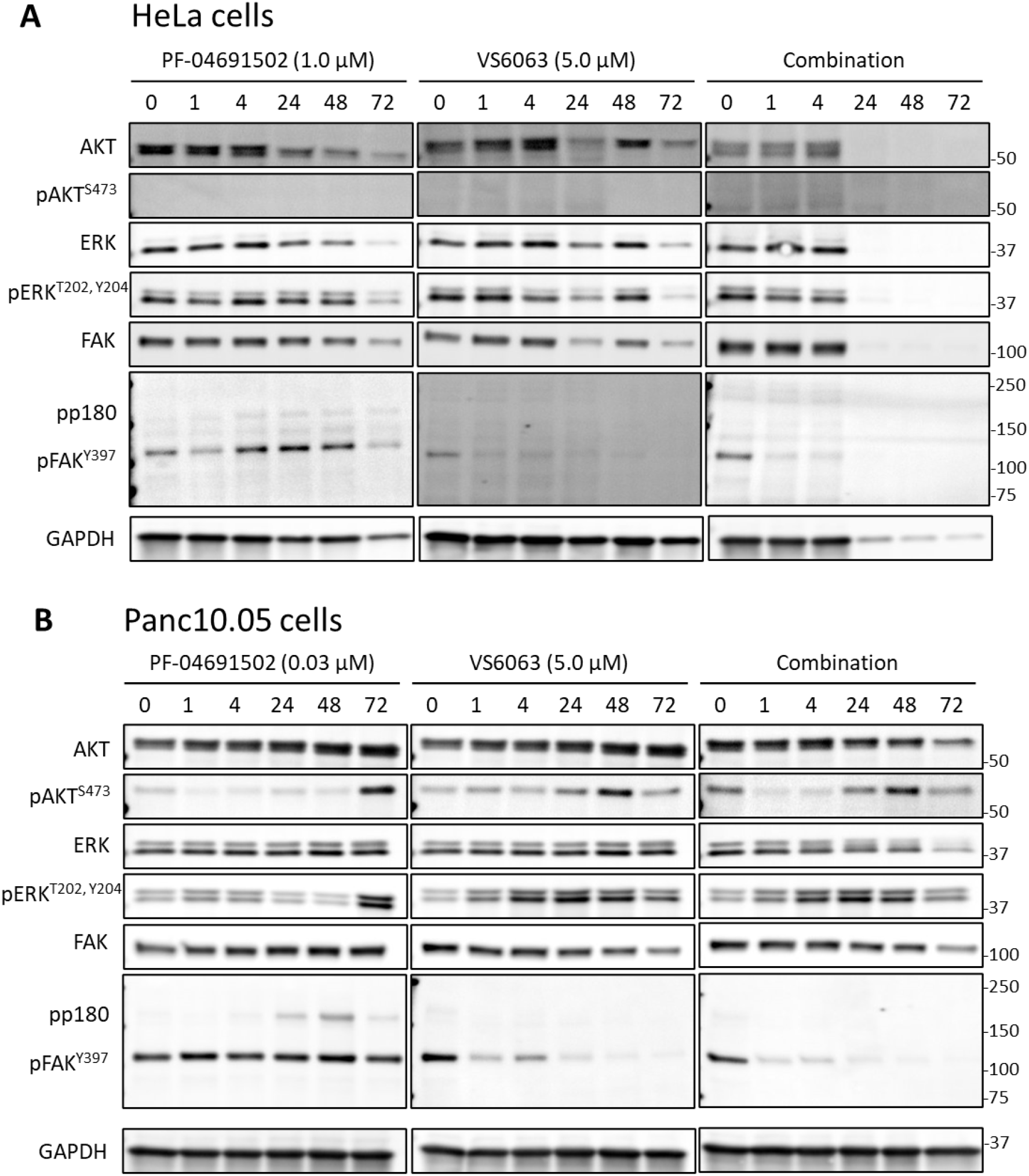
Immunoblotting of HeLa cells treated with PI3K and FAK inhibitors. HeLa cells (A) and Panc10.05 (B) cells were treated with PF-04691502, VS-6063, or the combination of the two inhibitors. Samples were taken at the indicated time points for immunoblotting of total and phosphorylated forms of AKT, FAK, and ERK, as well as the GAPDH loading control.

### PI3K inhibition induces activation of multiple RTKs

Interestingly, cells treated with PF-04691502 showed extra bands above the FAK band (MW: 125 kDa) after 24-48 hours in the phospho-FAK immunoblot (**Fig. 3A, B, red boxes**). The size of the bands varied among the cell lines, ranging from molecular weight of 150∼180 kD (designated pp180) except for SiHa, in which the size was smaller than 150 kD. Treating cells with ZSTK474, a pan-class I PI3K inhibitor, induced similar bands in a dose- and time-dependent manner (**Fig. 5A, B**). Since the phospho-FAK antibody is known to cross-react with tyrosine phosphorylated proteins such as EGFR, we tested whether pp180 represented activation of EGFR and HER2, which have similar molecular weights. We found that PF-04691502 treatment led to significant increase in the levels of EGFR, phospho-EGFR, HER2 and phospho-HER2 in pancreatic cancer cell lines Panc10.05 and A6L (**Fig. 5C**). Although EGFR and HER2 levels were also increased in cervical cancer cell lines, they were not accompanied by an increase in the phosphorylated forms of these proteins.

**Figure 5.**
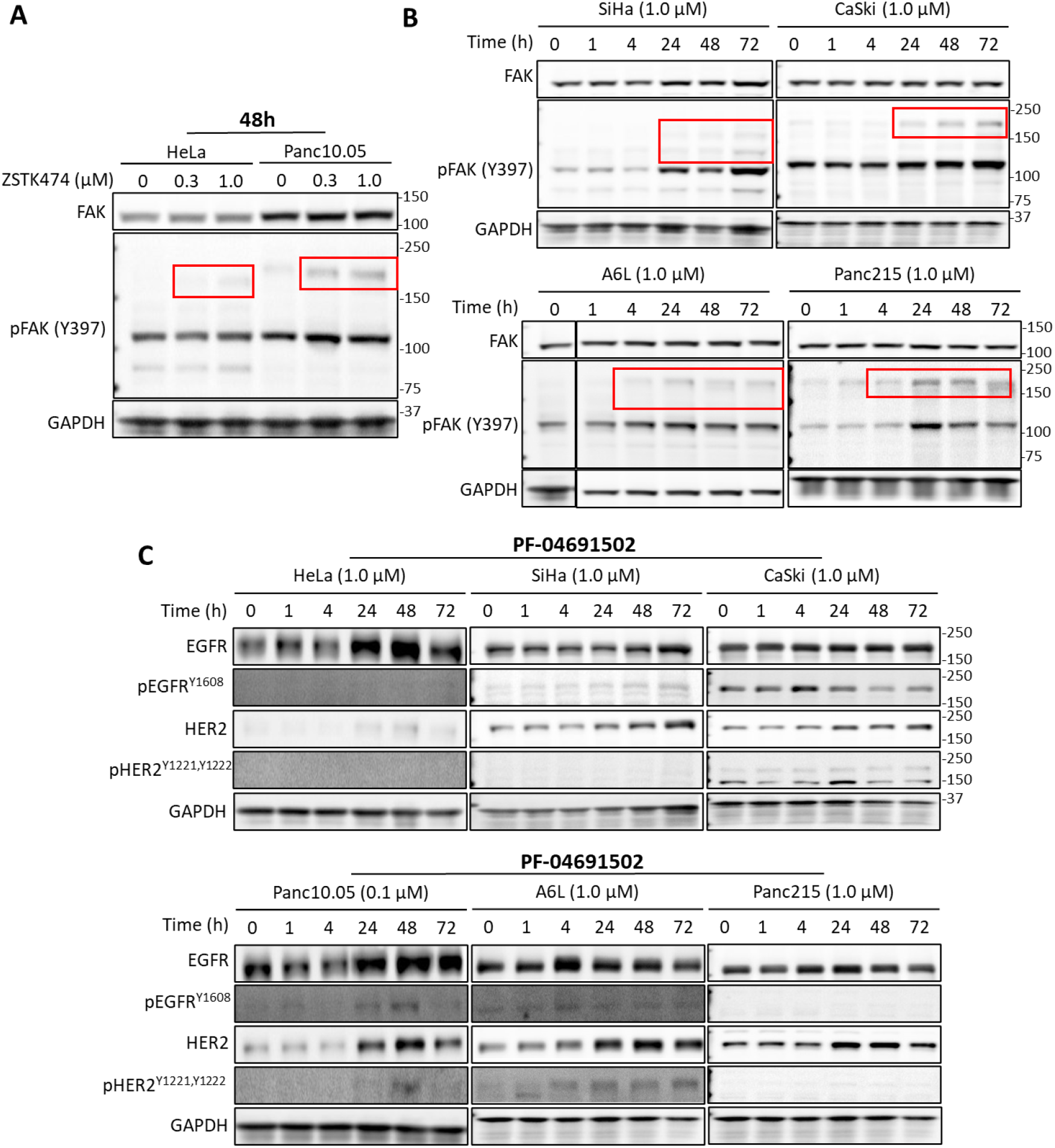
Immunoblotting of cells treated with PI3K inhibitors. (A-B) Immunoblots of FAK and phospho-FAK for HeLa and Panc10.05 cells treated with different concentrations of ZSTK474. (B) Immunoblots of FAK and phospho-FAK for cervical (top) and pancreatic (bottom) cancer cells treated with ZSTK474 for the indicated duration. Red boxes indicate the unknown bands found in phospho-FAK immunoblots. (C) Immunoblots of EGFR, phospho-EGFR, HER2, and phospho-HER2 for cervical (top) and pancreatic (bottom) cancer cells treated with PF-04691502 for the indicated duration.

To comprehensively identify the RTKs that may be activated by PI3K inhibition, we performed phospho-RTK arrays on cells before and after treatment with PI3Ki for 48 hours. In HeLa cells, PF-04691502 treatment led to an increased phosphorylation of insulin R and IGF-1R, as well as decreased phosphorylation of HGFR and Dtk (**Fig. 6A**). In Panc10.05 cells, PF-04691502 treatment led to an increased phosphorylation of EGFR, Her2, Her3, Axl, and EphA2 (**Fig. 6B**). Unlike the finding in HeLa cells, HGFR activity did not decrease in Panc10.05 after PF-04691502 treatment for 48h. An increased HER3 activity was also noticed in A6L cells after PF-04691502 treatment (**Fig. 6C**).

**Figure 6.**
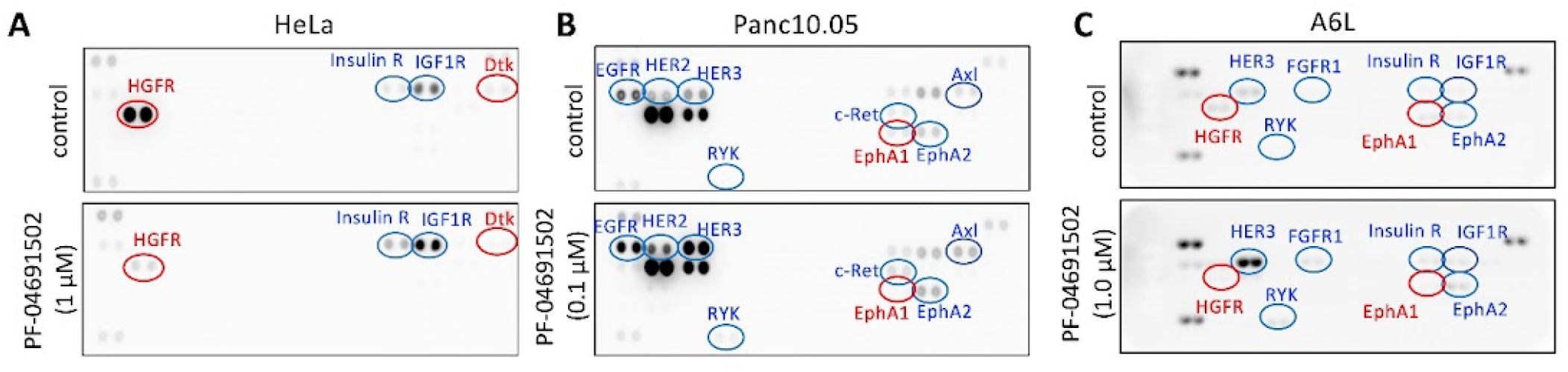
Phospho-RTK array of HeLa, Panc10.05, and A6L cells. Cells were treated with DMSO or PF-04691502 for 48 hours and the phospho-RTK activity was probed using the Proteome Profiler Human Phospho-RTK Array Kit (R&D Systems, ARY001B). Red and green ovals denote spots showing decreased and increased intensity, respectively, after PF-04691502 treatment.

## Discussion

In this study we explored the idea of targeting excitability of the Ras signaling network for cancer treatment. We previously identified a positive feedback between Ras, PI3K, the cytoskeleton, and FAK that is required for the excitability [23]. Due to the redundancy in signaling networks, we reasoned that effective blockade of excitability requires simultaneous inhibition of more than one node in the feedback loop. Vertical inhibition of the same pathway has been found to be more effective than blocking a single node in cancer treatment, as exemplified by the combination of BRAF and MEK inhibitors in BRAF mutant melanoma [40,41]. Here we chose to test the effects of simultaneous FAK and PI3K inhibition on the growth and signaling of cancer cells.

FAK is commonly overexpressed in various cancer types including cervical cancer cells [35,36,42], and FAK inhibition reduces cervical cancer xenograft growth and metastasis[43], suggesting that FAK is a promising cancer target. However, as a single agent, FAK inhibitors showed limited efficacy. FAK has been implicated in the resistance to targeted therapy, and combinations of FAK inhibitors with other inhibitors may enhance the efficacy [37,42]. PI3K signaling plays a crucial role in the growth, proliferation, and survival of cells, making it an appealing target for cancer treatment. However, when PI3K inhibitors are used alone in clinical trials, their effectiveness is limited due to resistance to PI3K inhibition. The narrow therapeutic range and frequent treatment-related side effects further limit the efficacy of PI3K inhibitors. Therefore, finding ways to maximize the usefulness of these agents in patient treatment remains a challenging task [44,45].

While extensive crosstalks exist between FAK and PI3K signaling pathways [46], studies on simultaneous targeting of PI3K and FAK inhibitors are scarce. An earlier study reported potential synergy between PI3K and FAK inhibitors in lung cancer cells with reduced PTEN [47]. Our findings showed the synergy of PI3K and FAK inhibitors in select cervical and pancreatic cancer cell lines by increasing apoptosis and decreasing cell cycle progression. Interestingly, PI3K inhibition induced the activation of multiple RTKs including insulin receptor and IGF-1R in cervical cancer cells, as well as EGFR, Her2, Her3, Axl, and EphA2 in pancreatic cancer cells.

Earlier studies showed that pharmacological inhibition of PI3K signaling often leads to the development of adaptive resistance [45], and one common mechanism involves FOXO-mediated transcription of RTKs, including HER3, EGFR, and INSR/IGFR1 [48–50]. Resistance may also occur due to RTK-ligands interaction from autocrine production by the tumor cells themselves, paracrine contribution from the tumor stroma, or systemic production [51,52]. To combat these responses, combining PI3K pathway inhibitors with RTK inhibitors or blocking antibodies has been suggested as a potential solution [53–55].

In conclusion, our findings have demonstrated a synergy between PI3K and FAK inhibitors in suppressing the growth of select cervical and pancreatic cancer cells. This revelation underscores the potential of combining these inhibitors as a therapeutic strategy against these malignancies. However, it is important to note that while the synergy was observed in some cancer cell types, it was not universal across all subtypes within these tumor types. Hence, there arises a critical need for identifying and establishing biomarkers of sensitivity that can aid in identifying patients who would benefit the most from this potent combination therapy. Such biomarkers would be invaluable in personalized medicine and maximize treatment efficacy.

## Acknowledgments

The authors would like to thank Laura Wood for pancreatic cancer cell lines Panc10.05, A6L, and Panc215, as well as TC Wu and Chien-Fu Hung labs for assistance with flow cytometry analysis. This work was supported by NIH grants R01GM136711 (to C.H.H.), Cervical Cancer SPORE P50CA098252 Career Development Award (to J.M.Y.) and Pilot Project Award (to C.H.H.), and the Sol Goldman Pancreatic Cancer Research Center (to C.H.H.).

## Supplemental information

**Figure S1.**
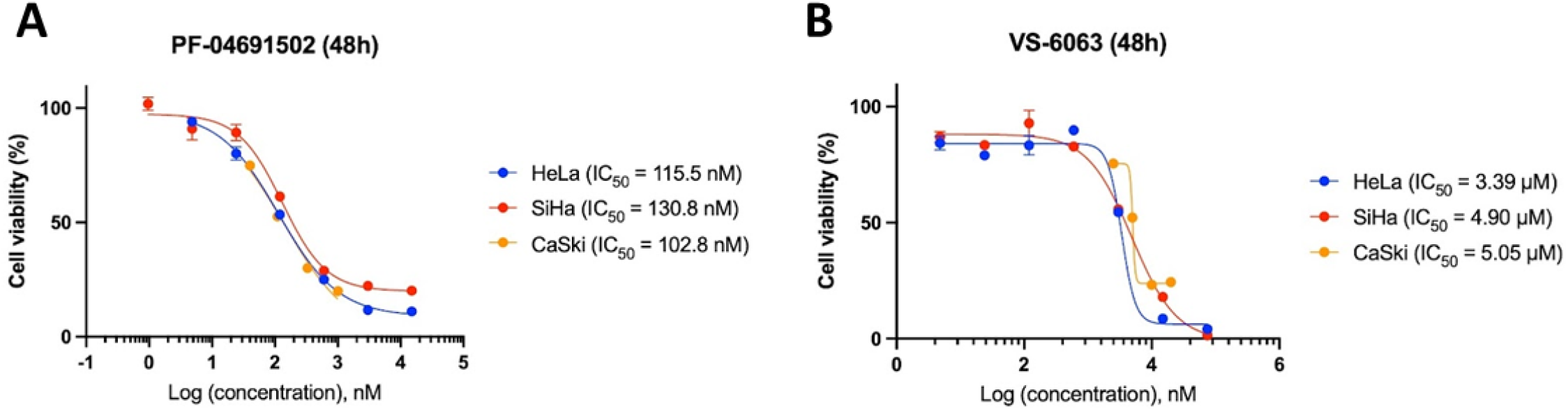
IC50 of PF-04691502 (A) and VS-6063 (B) in cervical cancer cell lines.

**Figure S2.**
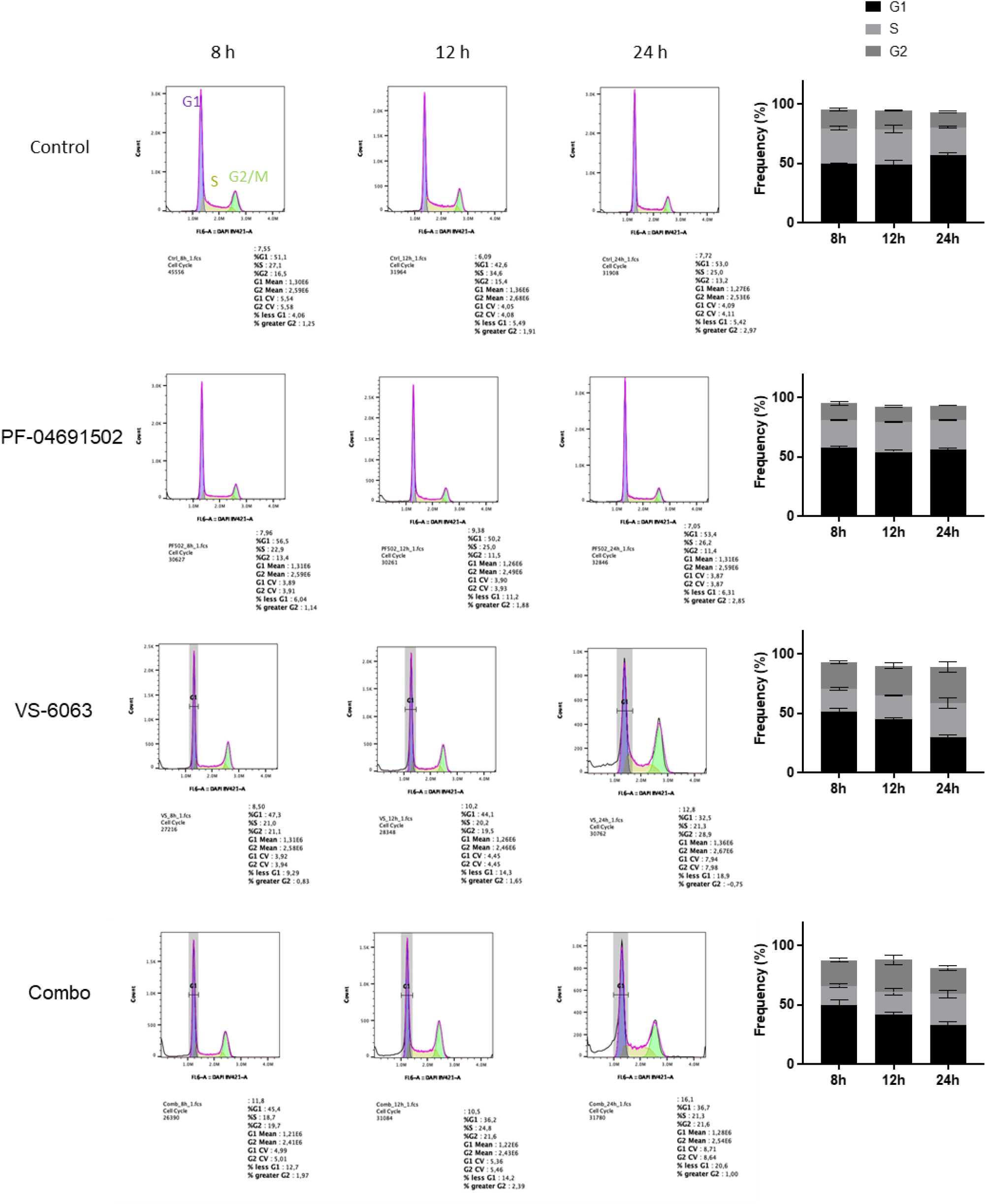
Cell cycle analysis of HeLa cells by flow cytometry of DAPI staining.

**Figure S3.**
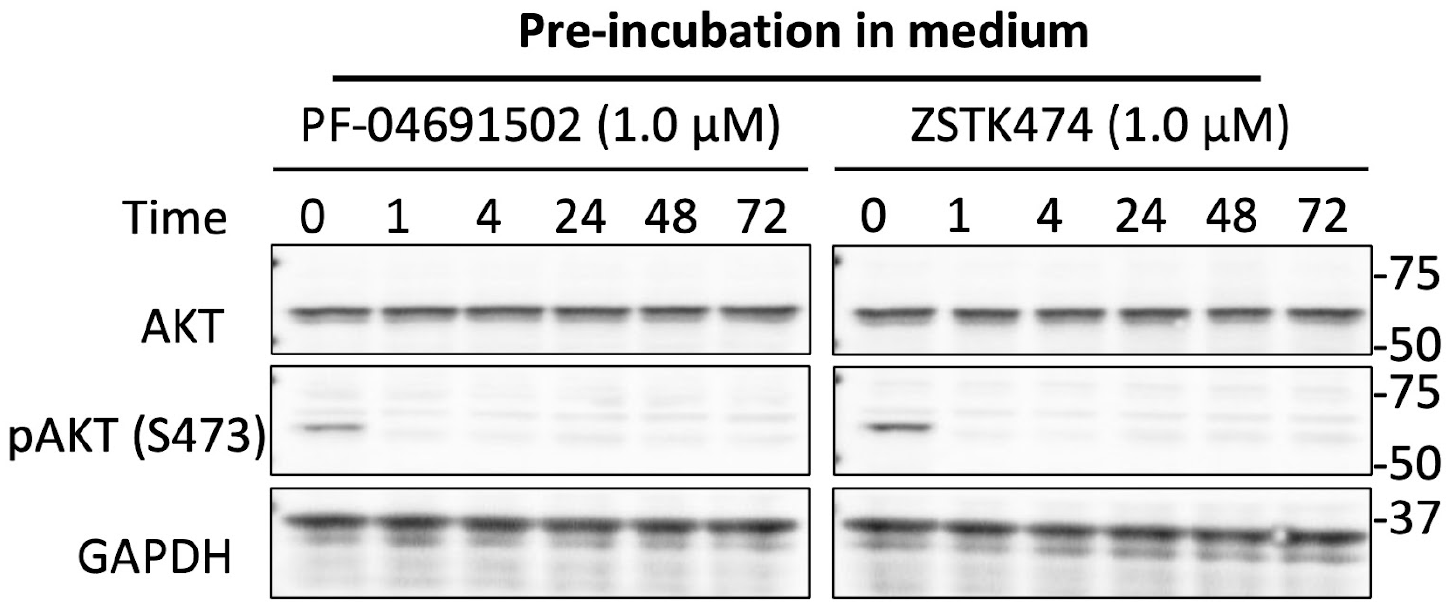
Stability of PI3K inhibitors. A6L cells were treated with the indicated inhibitors PF-04691502 or ZSTK474 for different periods. The medium was collected and transferred to another new plate of A6L cells. After 1 h incubation, cells were harvested for immunoblotting with the indicated antibodies.

**Figure S4.**
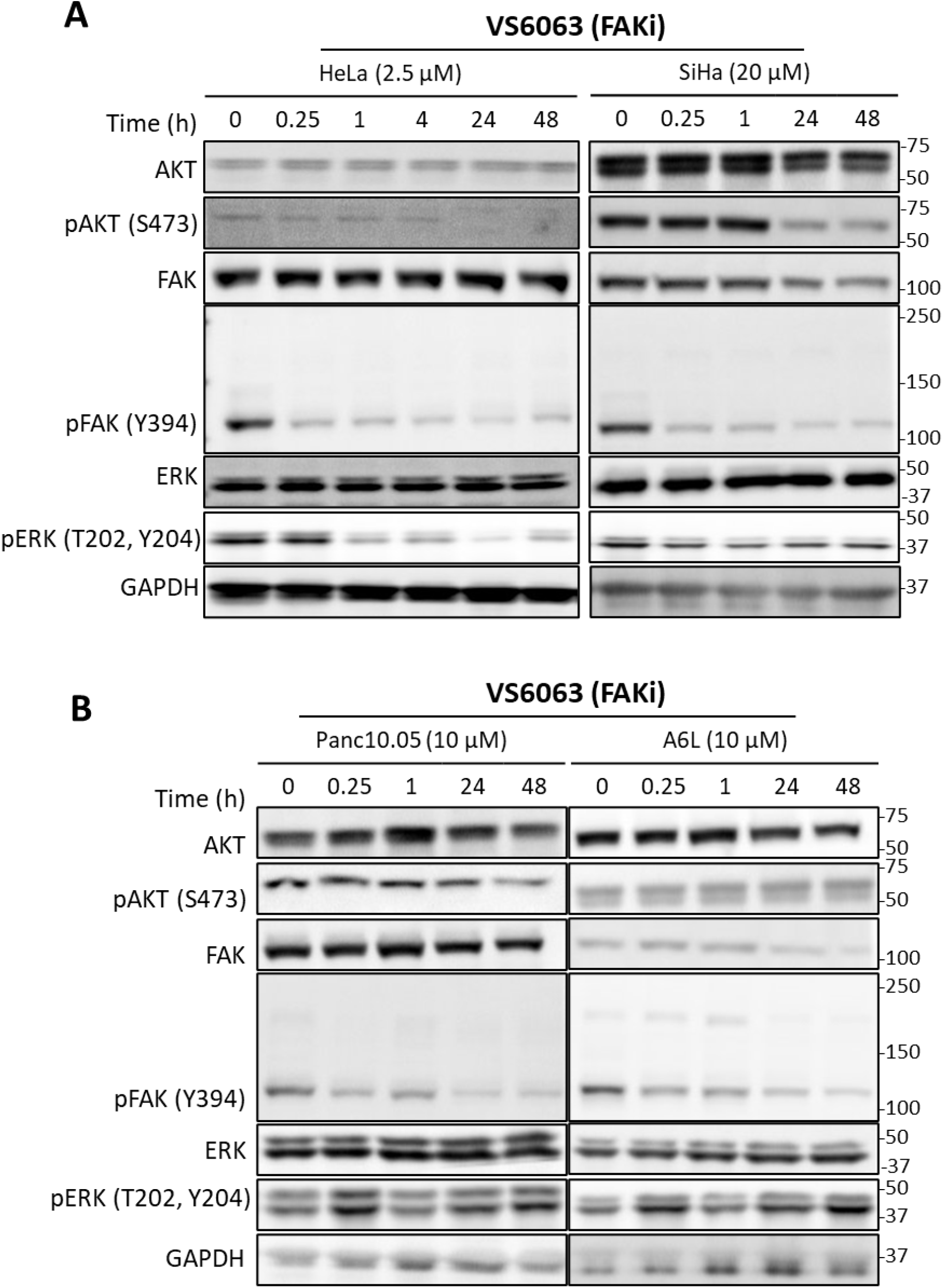
Immunoblotting of cervical and pancreatic cancer cells treated with FAK inhibitors. Cervical (A) and pancreatic (B) cancer cells were treated with VS-6063 and samples were taken at the indicated time points for immunoblotting of total and phosphorylated forms of AKT, FAK, and ERK, as well as the GAPDH loading control.

**Figure S5.**
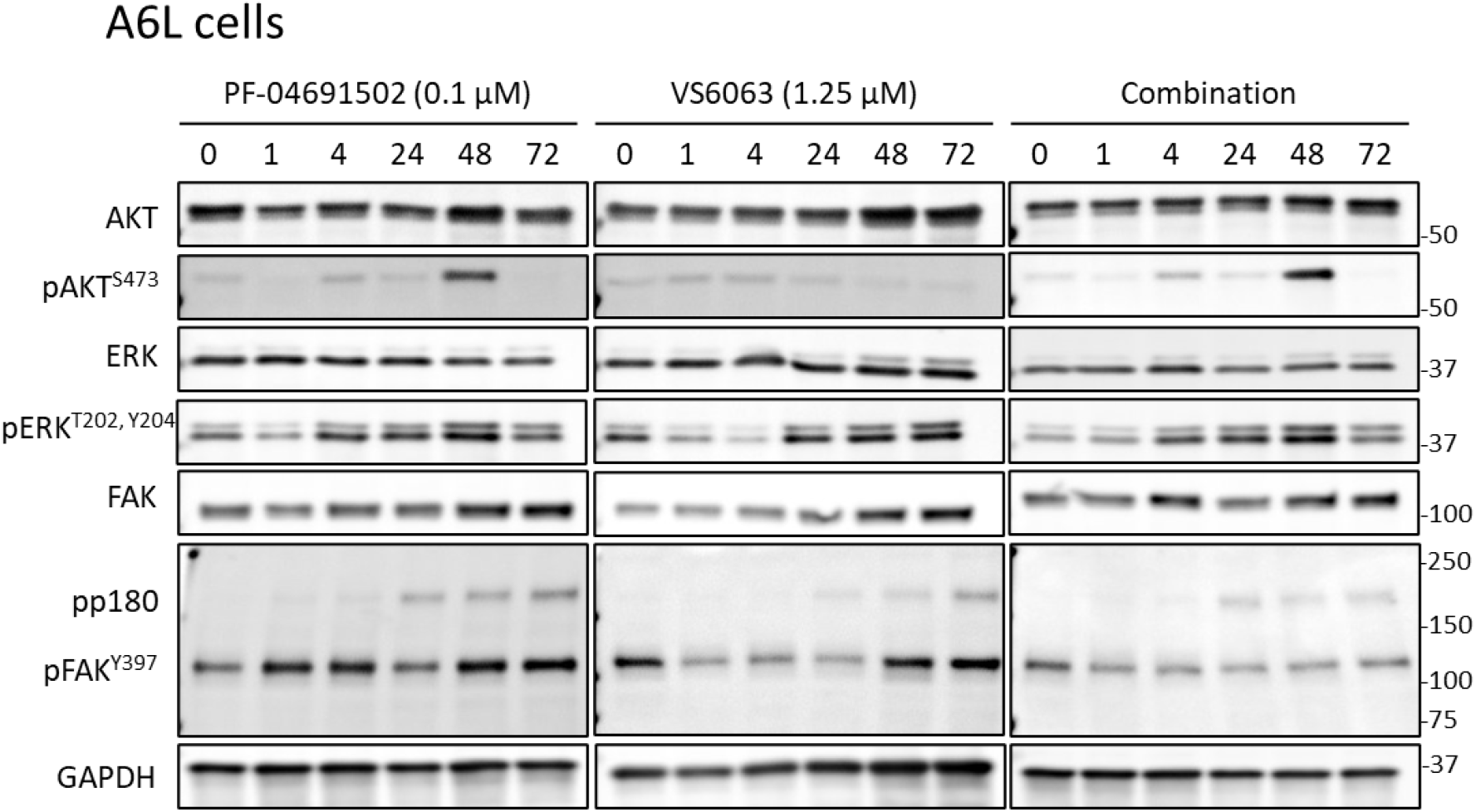
Immunoblotting of pancreatic cancer cell A6L treated with PI3K and FAK inhibitors. A6L cells were treated with PF-04691502, VS-6063, or the combination of the two inhibitors. Samples were taken at the indicated time points for immunoblotting of total and phosphorylated forms of AKT, FAK, and ERK, as well as the GAPDH loading control.

## References

1. Mendoza MC, Er EE, Blenis J. The Ras-ERK and PI3K-mTOR pathways: cross-talk and compensation. Trends Biochem Sci. 2011;36: 320–328. doi:10.1016/j.tibs.2011.03.006

2. Sanchez-Vega F, Mina M, Armenia J, Chatila WK, Luna A, La KC, et al. Oncogenic Signaling Pathways in The Cancer Genome Atlas. Cell. 2018;173: 321–337.e10. doi:10.1016/j.cell.2018.03.035

3. McIntyre JB, Wu JS, Craighead PS, Phan T, Köbel M, Lees-Miller SP, et al. PIK3CA mutational status and overall survival in patients with cervical cancer treated with radical chemoradiotherapy. Gynecol Oncol. 2013;128: 409–414. doi:10.1016/j.ygyno.2012.12.019

4. Ojesina AI, Lichtenstein L, Freeman SS, Pedamallu CS, Imaz-Rosshandler I, Pugh TJ, et al. Landscape of genomic alterations in cervical carcinomas. Nature. 2014;506: 371–375. doi:10.1038/nature12881

5. Miyake T, Yoshino K, Enomoto T, Takata T, Ugaki H, Kim A, et al. PIK3CA gene mutations and amplifications in uterine cancers, identified by methods that avoid confounding by PIK3CA pseudogene sequences. Cancer Lett. 2008;261: 120–126. doi:10.1016/j.canlet.2007.11.004

6. Janku F, Lee JJ, Tsimberidou AM, Hong DS, Naing A, Falchook GS, et al. PIK3CA mutations frequently coexist with RAS and BRAF mutations in patients with advanced cancers. PLoS One. 2011;6: e22769. doi:10.1371/journal.pone.0022769

7. Jiang W, Xiang L, Pei X, He T, Shen X, Wu X, et al. Mutational analysis of KRAS and its clinical implications in cervical cancer patients. J Gynecol Oncol. 2018;29: e4. doi:10.3802/jgo.2018.29.e4

8. Meira DD, de Almeida VH, Mororó JS, Nóbrega I, Bardella L, Silva RLA, et al. Combination of cetuximab with chemoradiation, trastuzumab or MAPK inhibitors: mechanisms of sensitisation of cervical cancer cells. Br J Cancer. 2009;101: 782–791. doi:10.1038/sj.bjc.6605216

9. Wright AA, Howitt BE, Myers AP, Dahlberg SE, Palescandolo E, Van Hummelen P, et al. Oncogenic mutations in cervical cancer: genomic differences between adenocarcinomas and squamous cell carcinomas of the cervix. Cancer. 2013;119: 3776–3783. doi:10.1002/cncr.28288

10. Hou M-M, Liu X, Wheler J, Naing A, Hong D, Coleman RL, et al. Targeted PI3K/AKT/mTOR therapy for metastatic carcinomas of the cervix: A phase I clinical experience. Oncotarget. 2014;5: 11168–11179. doi:10.18632/oncotarget.2584

11. Hou M-M, Liu X, Wheler J, Naing A, Hong D, Bodurka D, et al. Outcomes of patients with metastatic cervical cancer in a phase I clinical trials program. Anticancer Res. 2014;34: 2349–2355. Available: https://www.ncbi.nlm.nih.gov/pubmed/24778042

12. Cancer Genome Atlas Research Network. Electronic address, Cancer Genome Atlas Research Network. Integrated Genomic Characterization of Pancreatic Ductal Adenocarcinoma. Cancer Cell. 2017;32: 185–203.e13. doi:10.1016/j.ccell.2017.07.007

13. Waddell N, Pajic M, Patch A-M, Chang DK, Kassahn KS, Bailey P, et al. Whole genomes redefine the mutational landscape of pancreatic cancer. Nature. 2015;518: 495–501. doi:10.1038/nature14169

14. Nissim S, Leshchiner I, Mancias JD, Greenblatt MB, Maertens O, Cassa CA, et al. Mutations in RABL3 alter KRAS prenylation and are associated with hereditary pancreatic cancer. Nat Genet. 2019;51: 1308–1314. doi:10.1038/s41588-019-0475-y

15. Luchini C, Paolino G, Mattiolo P, Piredda ML, Cavaliere A, Gaule M, et al. KRAS wild-type pancreatic ductal adenocarcinoma: molecular pathology and therapeutic opportunities. J Exp Clin Cancer Res. 2020;39: 227. doi:10.1186/s13046-020-01732-6

16. Sasaki AT, Janetopoulos C, Lee S, Charest PG, Takeda K, Sundheimer LW, et al. G protein-independent Ras/PI3K/F-actin circuit regulates basic cell motility. J Cell Biol. 2007;178: 185–191. doi:10.1083/jcb.200611138

17. Asano Y, Nagasaki A, Uyeda TQP. Correlated waves of actin filaments and PIP3 in Dictyostelium cells. Cell Motil Cytoskeleton. 2008;65: 923–934. doi:10.1002/cm.20314

18. Weiger MC, Wang C-C, Krajcovic M, Melvin AT, Rhoden JJ, Haugh JM. Spontaneous phosphoinositide 3-kinase signaling dynamics drive spreading and random migration of fibroblasts. J Cell Sci. 2009;122: 313–323. doi:10.1242/jcs.037564

19. Arai Y, Shibata T, Matsuoka S, Sato MJ, Yanagida T, Ueda M. Self-organization of the phosphatidylinositol lipids signaling system for random cell migration. Proc Natl Acad Sci U S A. 2010;107: 12399–12404. doi:10.1073/pnas.0908278107

20. Taniguchi D, Ishihara S, Oonuki T, Honda-Kitahara M, Kaneko K, Sawai S. Phase geometries of two-dimensional excitable waves govern self-organized morphodynamics of amoeboid cells. Proc Natl Acad Sci U S A. 2013;110: 5016–5021. doi:10.1073/pnas.1218025110

21. Huang C-H, Tang M, Shi C, Iglesias PA, Devreotes PN. An excitable signal integrator couples to an idling cytoskeletal oscillator to drive cell migration. Nat Cell Biol. 2013;15: 1307–1316. doi:10.1038/ncb2859

22. Zhan H, Bhattacharya S, Cai H, Iglesias PA, Huang C-H, Devreotes PN. An Excitable Ras/PI3K/ERK Signaling Network Controls Migration and Oncogenic Transformation in Epithelial Cells. Dev Cell. 2020;0. doi:10.1016/j.devcel.2020.08.001

23. Yang J-M, Bhattacharya S, West-Foyle H, Hung C-F, Wu T-C, Iglesias PA, et al. Integrating chemical and mechanical signals through dynamic coupling between cellular protrusions and pulsed ERK activation. Nat Commun. 2018;9: 4673. doi:10.1038/s41467-018-07150-9

24. Yang J-M, Chi W-Y, Liang J, Takayanagi S, Iglesias PA, Huang C-H. Deciphering cell signaling networks with massively multiplexed biosensor barcoding. Cell. 2021;184: 6193–6206.e14. doi:10.1016/j.cell.2021.11.005

25. Albeck JG, Mills GB, Brugge JS. Frequency-modulated pulses of ERK activity transmit quantitative proliferation signals. Mol Cell. 2013;49: 249–261. doi:10.1016/j.molcel.2012.11.002

26. Aoki K, Kumagai Y, Sakurai A, Komatsu N, Fujita Y, Shionyu C, et al. Stochastic ERK activation induced by noise and cell-to-cell propagation regulates cell density-dependent proliferation. Mol Cell. 2013;52: 529–540. doi:10.1016/j.molcel.2013.09.015

27. Regot S, Hughey JJ, Bajar BT, Carrasco S, Covert MW. High-sensitivity measurements of multiple kinase activities in live single cells. Cell. 2014;157: 1724–1734. doi:10.1016/j.cell.2014.04.039

28. Aoki K, Kondo Y, Naoki H, Hiratsuka T, Itoh RE, Matsuda M. Propagating Wave of ERK Activation Orients Collective Cell Migration. Dev Cell. 2017;43: 305–317.e5. doi:10.1016/j.devcel.2017.10.016

29. Hino N, Rossetti L, Marín-Llauradó A, Aoki K, Trepat X, Matsuda M, et al. ERK-Mediated Mechanochemical Waves Direct Collective Cell Polarization. Dev. Cell. 2020. p. 2019.12.25.888552. doi:10.1016/j.devcel.2020.05.011

30. Hiratsuka T, Fujita Y, Naoki H, Aoki K, Kamioka Y, Matsuda M. Intercellular propagation of extracellular signal-regulated kinase activation revealed by in vivo imaging of mouse skin. Elife. 2015;4: e05178. doi:10.7554/eLife.05178

31. Tang M, Wang M, Shi C, Iglesias PA, Devreotes PN, Huang C-H. Evolutionarily conserved coupling of adaptive and excitable networks mediates eukaryotic chemotaxis. Nat Commun. 2014;5: 5175. doi:10.1038/ncomms6175

32. Nishikawa M, Hörning M, Ueda M, Shibata T. Excitable signal transduction induces both spontaneous and directional cell asymmetries in the phosphatidylinositol lipid signaling system for eukaryotic chemotaxis. Biophys J. 2014;106: 723–734. doi:10.1016/j.bpj.2013.12.023

33. Ryu H, Chung M, Dobrzyński M, Fey D, Blum Y, Lee SS, et al. Frequency modulation of ERK activation dynamics rewires cell fate. Mol Syst Biol. 2015;11: 838. doi:10.15252/msb.20156458

34. Ozkan-Dagliyan I, Diehl JN, George SD, Schaefer A, Papke B, Klotz-Noack K, et al. Low-Dose Vertical Inhibition of the RAF-MEK-ERK Cascade Causes Apoptotic Death of KRAS Mutant Cancers. Cell Rep. 2020;31: 107764. doi:10.1016/j.celrep.2020.107764

35. McCormack SJ, Brazinski SE, Moore JL Jr, Werness BA, Goldstein DJ. Activation of the focal adhesion kinase signal transduction pathway in cervical carcinoma cell lines and human genital epithelial cells immortalized with human papillomavirus type 18. Oncogene. 1997;15: 265–274. doi:10.1038/sj.onc.1201186

36. Golubovskaya VM. Focal adhesion kinase as a cancer therapy target. Anticancer Agents Med Chem. 2010;10: 735–741. Available: https://www.ncbi.nlm.nih.gov/pubmed/21214510

37. Dawson JC, Serrels A, Stupack DG, Schlaepfer DD, Frame MC. Targeting FAK in anticancer combination therapies. Nat Rev Cancer. 2021;21: 313–324. doi:10.1038/s41568-021-00340-6

38. Vanhaesebroeck B, Perry MWD, Brown JR, André F, Okkenhaug K. PI3K inhibitors are finally coming of age. Nat Rev Drug Discov. 2021; 1–29. doi:10.1038/s41573-021-00209-1

39. Ianevski A, Giri AK, Aittokallio T. SynergyFinder 2.0: visual analytics of multi-drug combination synergies. Nucleic Acids Res. 2020;48: W488–W493. doi:10.1093/nar/gkaa216

40. Long GV, Stroyakovskiy D, Gogas H, Levchenko E, de Braud F, Larkin J, et al. Combined BRAF and MEK inhibition versus BRAF inhibition alone in melanoma. N Engl J Med. 2014;371: 1877–1888. doi:10.1056/NEJMoa1406037

41. Flaherty KT, Infante JR, Daud A, Gonzalez R, Kefford RF, Sosman J, et al. Combined BRAF and MEK inhibition in melanoma with BRAF V600 mutations. N Engl J Med. 2012;367: 1694–1703. doi:10.1056/NEJMoa1210093

42. Sulzmaier FJ, Jean C, Schlaepfer DD. FAK in cancer: mechanistic findings and clinical applications. Nat Rev Cancer. 2014;14: 598–610. doi:10.1038/nrc3792

43. Schwock J, Dhani N, Cao MP-J, Zheng J, Clarkson R, Radulovich N, et al. Targeting focal adhesion kinase with dominant-negative FRNK or Hsp90 inhibitor 17-DMAG suppresses tumor growth and metastasis of SiHa cervical xenografts. Cancer Res. 2009;69: 4750–4759. doi:10.1158/0008-5472.CAN-09-0454

44. Yang J, Nie J, Ma X, Wei Y, Peng Y, Wei X. Targeting PI3K in cancer: mechanisms and advances in clinical trials. Mol Cancer. 2019;18: 26. doi:10.1186/s12943-019-0954-x

45. Fruman DA, Chiu H, Hopkins BD, Bagrodia S, Cantley LC, Abraham RT. The PI3K Pathway in Human Disease. Cell. 2017;170: 605–635. doi:10.1016/j.cell.2017.07.029

46. Zhao J, Guan J-L. Signal transduction by focal adhesion kinase in cancer. Cancer Metastasis Rev. 2009;28: 35–49. doi:10.1007/s10555-008-9165-4

47. Cavazzoni A, La Monica S, Alfieri R, Ravelli A, Van Der Steen N, Sciarrillo R, et al. Enhanced efficacy of AKT and FAK kinase combined inhibition in squamous cell lung carcinomas with stable reduction in PTEN. Oncotarget. 2017;8: 53068–53083. doi:10.18632/oncotarget.18087

48. Chakrabarty A, Sánchez V, Kuba MG, Rinehart C, Arteaga CL. Feedback upregulation of HER3 (ErbB3) expression and activity attenuates antitumor effect of PI3K inhibitors. Proc Natl Acad Sci U S A. 2012;109: 2718–2723. doi:10.1073/pnas.1018001108

49. Chandarlapaty S, Sawai A, Scaltriti M, Rodrik-Outmezguine V, Grbovic-Huezo O, Serra V, et al. AKT inhibition relieves feedback suppression of receptor tyrosine kinase expression and activity. Cancer Cell. 2011;19: 58–71. doi:10.1016/j.ccr.2010.10.031

50. Muranen T, Selfors LM, Worster DT, Iwanicki MP, Song L, Morales FC, et al. Inhibition of PI3K/mTOR leads to adaptive resistance in matrix-attached cancer cells. Cancer Cell. 2012;21: 227–239. doi:10.1016/j.ccr.2011.12.024

51. Zhang W, Huang P. Cancer-stromal interactions: role in cell survival, metabolism and drug sensitivity. Cancer Biol Ther. 2011;11: 150–156. doi:10.4161/cbt.11.2.14623

52. Wilson TR, Fridlyand J, Yan Y, Penuel E, Burton L, Chan E, et al. Widespread potential for growth-factor-driven resistance to anticancer kinase inhibitors. Nature. 2012;487: 505–509. doi:10.1038/nature11249

53. García-García C, Ibrahim YH, Serra V, Calvo MT, Guzmán M, Grueso J, et al. Dual mTORC1/2 and HER2 blockade results in antitumor activity in preclinical models of breast cancer resistant to anti-HER2 therapy. Clin Cancer Res. 2012;18: 2603–2612. doi:10.1158/1078-0432.CCR-11-2750

54. Garrett JT, Sutton CR, Kurupi R, Bialucha CU, Ettenberg SA, Collins SD, et al. Combination of antibody that inhibits ligand-independent HER3 dimerization and a p110α inhibitor potently blocks PI3K signaling and growth of HER2+ breast cancers. Cancer Res. 2013;73: 6013–6023. doi:10.1158/0008-5472.CAN-13-1191

55. Meister KS, Godse NR, Khan NI, Hedberg ML, Kemp C, Kulkarni S, et al. HER3 targeting potentiates growth suppressive effects of the PI3K inhibitor BYL719 in pre-clinical models of head and neck squamous cell carcinoma. Sci Rep. 2019;9: 9130. doi:10.1038/s41598-019-45589-y

